# Building synthetic cell-cell synapses using engineered scaffold protein condensates

**DOI:** 10.64898/2026.10.26.762062

**Authors:** Nick Y. Shin, Gaurav Mitra, Ania M. Grodsky, Kiersten M. Ruff, Gaurav Chauhan, Rohit V. Pappu, Wendell A. Lim

## Abstract

Synaptic assemblies that form at cell-cell junctions are often organized by multivalent scaffold proteins with condensate forming properties. An emerging hypothesis is that these scaffolds provide a thresholding mechanism whereby small increases in the density of cell-cell recognition receptors can trigger all-or-none assembly selectively at the correct partner cell interface. We test this hypothesis by engineering synthetic scaffolded synapses. Coarse-grained simulations suggest that properly-tuned scaffold interactions can yield a sharp condensation threshold. Guided by these simulations, we designed modular scaffold proteins consisting of PDZ domains and intrinsically disordered regions (IDRs) of different molecular grammars. Within a specific region of parameter space, controlled by valency, receptor binding affinity and IDR-driven condensation, the synthetic scaffold proteins exhibit switch-like condensation behaviors, where co-condensates of receptor and scaffold selectively form only at interfaces with the proper partner cell. Notably, orthogonal receptor-scaffold pairs enable a single receiver cell to form multiple distinct interfaces, each associated with a different partner cell. We demonstrate that these synthetic synapse assemblies can recruit different client proteins to enhance intercellular signaling, orient cell division, or sculpt complex synaptic morphologies. Our results illustrate how the ultrasensitive condensation properties of modular scaffold proteins can be harnessed to create novel and highly specific cell-cell interfaces.

## Introduction

Multicellular organisms rely on specialized cell–cell synapses to transmit information and coordinate tissue-level behavior (**Fig. 1a**).^1,2^ These cell junction assemblies are critical for neuronal wiring,^3^ formation of epithelial junctions,^4^ and immune surveillance.^5^ A fundamental question is how cells form these concentrated, membrane-associated assemblies only at interfaces with the correct partner cell.^2,3,5^

**Figure 1.**
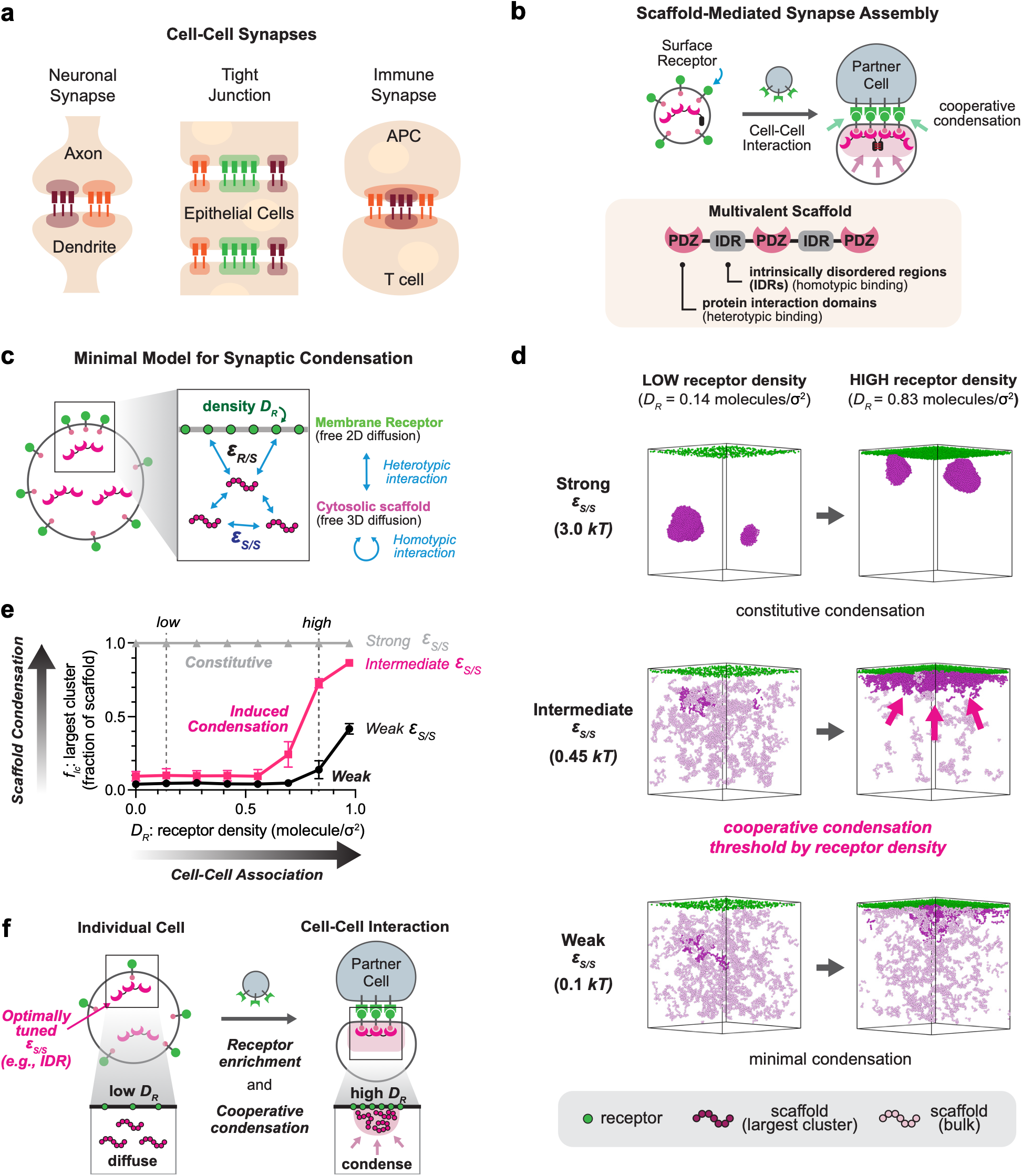
Multivalent scaffold proteins provide a general physical mechanism for cell contact-induced synapse assembly. a, Diverse cell-cell synapses, including neuronal, tight junction, and immune synapses, rely on scaffold proteins to concentrate signaling proteins at sites of contact. b, Conceptual model of scaffold-mediated synapse assembly. Cell-to-cell interactions cluster surface receptors, driving the local cooperative condensation of multivalent cytosolic scaffolds containing intrinsically disordered regions (IDRs) and interacting domains. c, Coarse-grained (CG) molecular dynamics simulation model of synaptic condensation. Receptors are can diffuse laterally in a two-dimensional membrane plane on the upper surface of the simulation box, representative of the membrane. and Scaffolds molecules (which are represented by 25-bead homopolymeric chains in our CG simulations) can freely diffuse in an adjacent three-dimensional volume. Receptor-scaffold interaction strength ε_R/S_ is fixed at 15 kT; receptor density D_R_ and scaffold–scaffold interaction strength ε_S/S_ are varied. The simulations were performed using LAMMPS. d, Representative simulation snapshots of the final frame showing scaffold condensation with varied scaffold-scaffold self-interaction strengths (ε_S/S_) and receptor densities (D_R_ of 0.14 and 0.83 molecules/σ²). Receptors are shown in green, scaffold molecules that belong to the largest connected cluster are shown in dark purple, and the rest of the scaffolds molecules are shown in light purple. Here, σ is the diameter of each receptor in our CG simulations. The area of the 2D plane representing the membrane is (60σ)^2^ = 3600σ^2^. A receptor density D_R_=1, then corresponds to 3600 receptors. e, Fraction of scaffold molecules in the largest connected cluster (f_lc_) as a function of receptor density (D_R_). Weak self-interaction (0.10 kT) produces little assembly, intermediate self-interaction (0.45 kT) produces receptor-density-dependent assembly, and strong self-interaction (3.0 kT) produces constitutive condensation. The error bars represent the standard deviation, computed across 5 simulation replicates and the final 10 frames of the fully converged simulations. f, Proposed biophysical model. Properly tuned scaffold–scaffold interactions prevent spontaneous phase separation in isolated cells but trigger cooperative condensation upon contact-induced receptor enrichment at the cell-cell interface.

A recurring feature of cell-cell synapses is their organization by multivalent scaffold proteins. Postsynaptic density scaffolds such as PSD-95 and related MAGUK proteins,^6,7^ epithelial junctional scaffolds such as ZO proteins,^8,9^ and immune-signaling adaptors such as LAT and SLP-76 share a general modular architecture consisting of multiple protein interaction domains and intrinsically disordered regions (IDRs).^5,10–13^ The IDRs function either as linkers or terminal tail domains.^14,15^ Importantly, many scaffold proteins also bind client proteins, such as receptors and downstream effectors, recruiting them to the synaptic complex. This general design of scaffold proteins has deep evolutionary roots: early animals such as sponges have conserved MAGUK/PDZ scaffolds, homologous to those that were later used at neuronal synapses.^16,17^ Thus, scaffold proteins appear to have first evolved as organizers of fundamental intercellular communication within early multi-celled organisms, predating the evolution of neurons.^11,17^

A growing body of work suggests that biomolecular condensation is a key physical mechanism by which scaffold proteins organize synaptic complexes (**Fig. 1b**).^18–25^ Condensates form via phase transitions of multivalent proteins comprising folded domains and IDRs. These transitions combine phase separation, a density transition, and percolation, a networking transition.^14,19,26^ These observations have led to the hypothesis that multivalent scaffold proteins with condensation properties may provide a thresholding mechanism by which small increases in the local density of receptors that recognize cognate partner cells could conditionally induce the formation of synaptic assemblies.^24,27,28^ This hypothesis is consistent with recent findings where multivalent interaction networks and their condensates were found to drive ultra sensitivity across diverse pathways, such as Arp2/3-mediated actin polymerization, Ras activation in TCR signaling clusters, and transcriptional coactivators at super-enhancers.^19,20,29^ Biomolecular condensation may therefore act as an ultrasensitive process that translates subtle shifts in physiological input into sharp, non-linear outputs, similar to how correct partner cell contact triggers synaptic organization.^18,30,31^

Thus, we reasoned that a modular scaffold design comprising repeated receptor binding domains and self-associating IDRs might be sufficient to generate minimal synthetic cell-cell synapses.^32^ We use the term synapse in an operational, architectural sense to refer to a defined molecular complex that forms at a partner-selective cell–cell contact.^27^ Here we first test the plausibility of this design through coarse-grained molecular dynamics simulation and use the resulting principles to guide the engineering of synthetic cell-cell synapses in mammalian cells.

### Simulations define scaffold parameter regimes that can yield receptor density dependent condensation

When two partner cells first bind each other through receptors, these receptors will become locally focused at the contact zone and undergo a local increase in density.^1^ We used simulations to explore the plausibility that a small increase in local receptor density could drive switch-like scaffold condensation, which is the pre-requisite for partner-selective condensation-mediated synapse organization. We considered a reduced model in which cytoplasmic scaffold molecules engage in homotypic interactions with one another and heterotypic interactions with surface receptors (**Fig. 1c**). We built a coarse-grained molecular dynamics model using LAMMPS (Large-scale Atomic/Molecular Massively Parallel Simulator) ^33^ (see Methods for simulation details). The model contains laterally mobile membrane receptors (represented by individual beads) confined to a two-dimensional plane on the surface, and scaffold molecules diffusing in an adjacent three-dimensional volume. The homotypic interactions between polymer beads are designated as ε_S/S_. Multivalence of cohesive motifs is captured by the number of beads in the polymer capable of encoding equivalent inter-bead interactions of strength ε_S/S_. We chose 25 beads to mimic the number of cohesive motifs known as stickers that are observed across homologs of prion-like low complexity domains.^32,34^

Receptor beads bind scaffold beads with interaction strength ε_R/S_. We fixed the heterotypic receptor–scaffold interaction strength at 15 *kT*, which is representative of sub-micromolar binding affinity typical of weak protein binding domains such as SH3 and PDZ.^35–37^ We then varied scaffold–scaffold attraction and receptor surface density, D_R_. In this formulation, receptor density is a control parameter that models cell-cell binding: low D_R_ represents the dispersed pre-contact state, whereas high D_R_ approximates receptor focusing on an engaged cell–cell interface.

We first determined how scaffold self-attraction controls bulk condensation in the near-absence of receptors (D_R_ = 0.0003 molecules/σ^2^, σ: diameter of receptor beads). As a metric for bulk condensation we measured f_lc_ – the fraction of scaffold molecules contained in the largest cluster. The model exhibited a sharp sigmoidal increase in f_lc_, at ε_S/S_ of approximately 0.4873 kT (**Extended Data Fig. 1a, 1b**), beyond which bulk condensation is constitutively observed. We therefore selected ε_S/S_ = 0.45 kT, just below this bulk condensation threshold, as the intermediate regime of scaffold-scaffold self-association. Together with the strong and weak regime (ε_S/S_ of 3.0 kT and 0.1 kT, respectively), we asked whether increasing receptor density could trigger localized assembly at the membrane (**Fig. 1d**). The results demonstrate the following: Strong scaffold homotypic interaction (ε_S/S_ = 3.0 kT) produces constitutive scaffold condensation with little dependence on the receptor density, while weak homotypic interaction (ε_S/S_ = 0.10 kT) produces little assembly even at high receptor density. In contrast, inducible, receptor density-dependent condensation of scaffolds is possible when the homotypic associations of the scaffold are in an intermediate regime. When the scaffold self-association is maintained at the intermediate regime (ε_S/S_ = 0.45 kT), scaffold molecules remain largely diffuse in the cytoplasm at low receptor density (D_R_ = 0.14 molecules/σ²). However, they preferentially accumulate near the membrane at a high receptor density (D_R_ = 0.83 molecules/σ²). Here, the order parameter f_lc_ shows non-linear, sigmoidal dependence on the receptor density, representing ultra sensitivity (**Fig. 1e**).

What emerges is a physical model with the following features: when scaffold self-interaction strength is in an appropriate, albeit intermediate level, the local receptor clustering upon cell-cell interaction can be sufficient to yield switch-like membrane-localized condensation (**Fig. 1f**). Thus, the simple model supports the idea that a condensing scaffold has the potential to drive highly interface selective assembly at a membrane when the interactions are tuned to ensure that scaffold self-associations are neither too strong (constitutive condensation), nor too weak (no condensation), but are just right to be coupled to receptor binding to enable responsive condensation. In real systems, the tunability of self-associations can be achieved through the design of molecular grammars of IDRs, the choice of modular protein-protein interaction domains, and the ligands to which these domains bind.^19,20,38,39^

### Engineered scaffolds reconstitute partner cell-induced condensate formation

We next asked whether the predicted regime of cell contact-induced scaffold condensation could be realized using genetically encoded components in living cells (here L929 fibroblast cells). We engineered receiver cells to express a synthetic surface receptor containing extracellular GFP and an intracellular PDZ-binding ligand. We also engineered cognate partner cells that express membrane tethered αGFP nanobodies on their surface (**Fig. 2a**).^40^ Partner cell engagement can thereby concentrate the GFP-fused receptor at the cell–cell interface.

**Figure 2.**
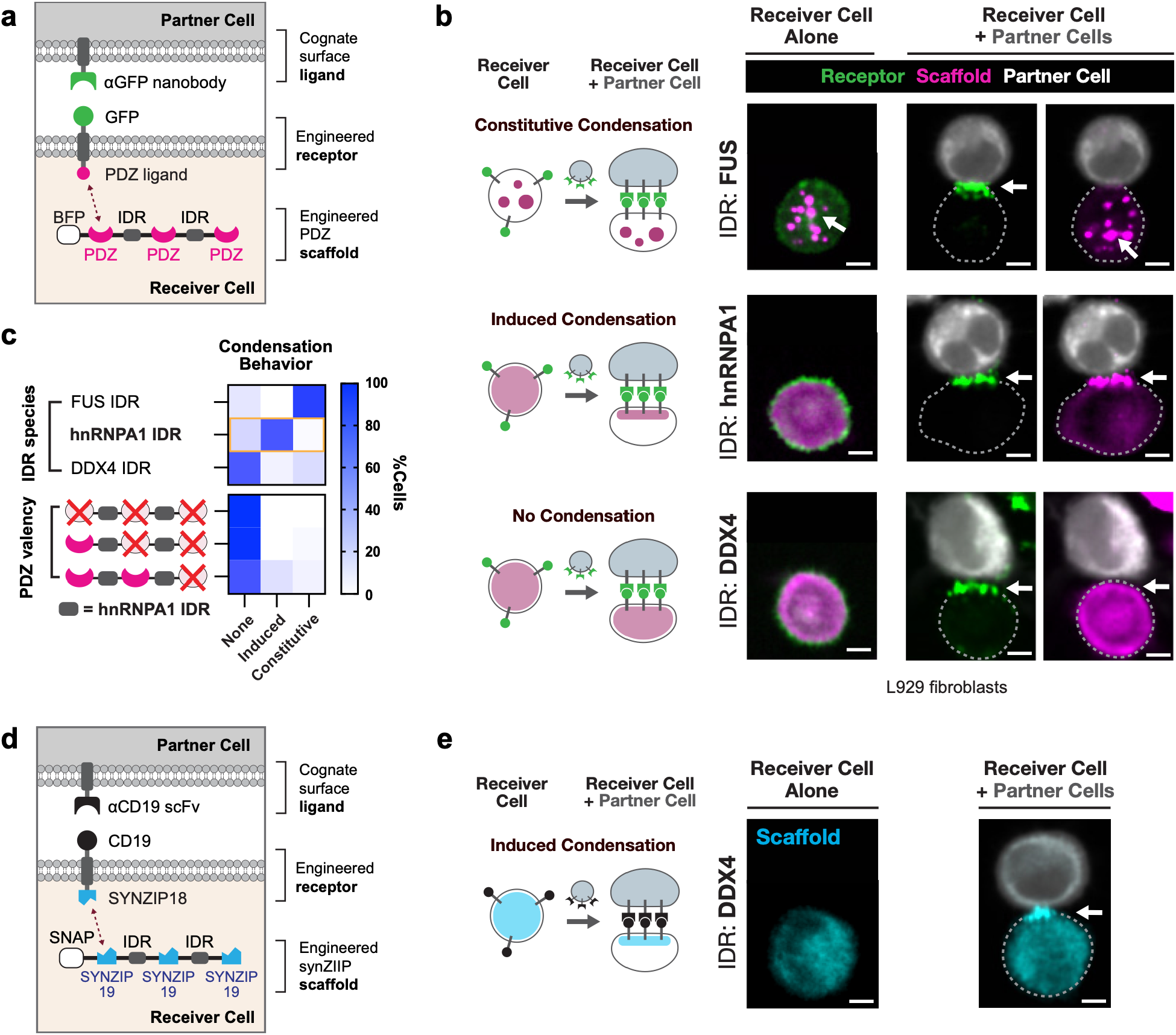
Engineered multivalent scaffolds drive contact-induced synaptic condensation in living cells. a, Construct design of the engineered cell-cell interface. A surface ligand on the partner cell (αGFP nanobody) engages a cognate GFP receptor on the receiver cell, recruiting cytosolic multivalent scaffolds containing PDZ interaction domains connected by intrinsically disordered regions (IDRs). b, Representative confocal images of L929 receiver cells expressing PDZ scaffolds with the indicated IDRs, cultured alone or with αGFP partner cells. Receptor is shown in green, scaffold in magenta and partner cells in white (labelled with CellTrace Far Red). Scale bars, 5 µm; images were acquired 2 h after co-culture. c, Combinatorial matrix depicting scaffold condensation state (%cells showing none, cell contact-induced, or constitutive condensation) as a function of IDRs and PDZ domain valency. (n = 59, 60, and 60 for FUS IDR, hnRNPA1 IDR, and DDX4 IDR, respectively, and n = 63, 73, and 72, for PDZ valency of 0, 1, and 2, respectively.) Matched scaffold-expression measurements are shown in Extended Data Fig. 3d. d, Construct design of an orthogonal CD19/αCD19 interface using a SYNZIP receptor–scaffold interaction. e, Representative confocal images showing contact-induced assembly of a DDX4 IDR-containing SYNZIP scaffold at cognate interfaces. Scaffold is shown in cyan, and partner cells in white (labelled with CellTrace Far Red). Scale bars, 5 µm; 2 h after co-culture. Quantification and expression matching are shown in Extended Data Fig. 5.

To couple receptor enrichment to cooperative scaffold assembly, we designed a multivalent cytoplasmic scaffold protein analogous to MAGUK-family scaffold proteins (**Extended Data Fig. 2a**). PSD-95, ZO-1 and Dlg1 exemplify the modular architecture of MAGUK-family scaffolds, where folded binding domains (e.g., PDZ and SH3) are interspersed with IDRs that can generate multivalent networks while keeping individual interactions relatively weak. We then performed a bioinformatics survey of human proteome containing long IDRs and cross-referenced it with cytoplasmic cell-cell junction proteins (**Extended Data Fig. 2b**, see Methods). The results highlight a recurring combination of repeated protein–interaction domains and multiple disordered segments. Notably, when we focus on PDZ domain-containing proteins, they most commonly possess three protein-binding domains and two distinct IDRs. We therefore built a minimal scaffold containing three PDZ interaction domains separated by two IDRs. To minimize any cross-reaction of our engineered proteins with native proteins, we used engineered Erbin PDZ mutants and their cognate ligands, as these variants are orthogonal to the major classes of endogenous PDZ domains (see Methods).^41^ For visualization, each scaffold construct was fused to BFP at the N-terminus.

We then selected low-complexity IDRs from hnRNPA1, FUS and DDX4, proteins whose disordered regions have well-characterized condensation propensities.^32,42–44^ In L929 fibroblasts, receiver cells were introduced with the GFP-fused receptor and PDZ scaffold proteins containing each IDR. The basal state of condensation was measured by confocal microscopy. Then, the receiver cells were co-cultured with the αGFP partner cells for 2 hours, followed by confocal microscopy imaging. We observed that the hnRNPA1 IDR-containing scaffold showed the target behavior of partner cell-induced synaptic condensation. Specifically, the scaffold was diffuse in receiver cells cultured alone but became strongly enriched in punctate assemblies at interfaces with αGFP partner cells (**Fig. 2b**). The assembly was sharply restricted to sites of partner engagement, consistent with the cell contact-induced condensation predicted from coarse-grained molecular dynamics simulations.

Changing the IDR sequence, however, shifted the scaffold into different assembly regimes. The scaffold construct containing N-terminal prion-like low complexity domain (PLCD) of FUS (hereby FUS IDR) formed cytoplasmic condensates even in the absence of partner cells, corresponding qualitatively to the constitutive regime predicted for strong homotypic attractions. By contrast, the DDX4 IDR-containing scaffold remained largely diffuse before and after partner cell engagement, never forming condensates, even with partner cell engagement. Based on these results and the simulations, we classified the scaffold condensation behaviors into three distinct regimes; constitutive condensation, where the scaffold puncta form spontaneously before partner cell contact; induced condensation, where scaffold puncta are only observed at the interface against the partner cell; and no condensation, where scaffold remains diffuse in all conditions. When the single-cell condensation behaviors were quantified as percentage of cells occupying each condensation regime, the results were consistent with the representative confocal microscopy images (**Fig. 2c**).

To test the effect of IDR self-attraction, we replaced IDRs with a flexible (GGGGS)_3_ linker or mutants of hnRNPA1 IDR (**Extended Data Fig. 3a, c**).^32^ Phase separation coupled to percolation of PLCDs from hnRNPA1 can be tuned by the valence, patterning, and ratio of Phe-to-Tyr residues with the higher Tyr content leading to stronger driving forces for condensation. Thus, changing the number of Tyr residues provides a handle for modulating homotypic interactions.^32^ Replacing the IDR with a flexible (GGGGS)_3_ linker keeps the scaffold diffuse before and after the partner cell engagement. When all 12 Tyr residues from hnRNPA1 IDR are replaced with Phe (hnRNPA1 IDR^YtoF^), thus significantly weakening its self-interaction, the scaffold remained largely diffuse. On the other hand, the hnRNPA1 IDR mutant with all 15 Phe residues replaced with Tyr (hnRNPA1 IDR^FtoY^) forms cytoplasmic condensates even in the absence of partner cells, as predicted from the strengthened IDR-IDR homotypic interaction. These results support the prediction that a proper tuning of IDR molecular grammars is required for cell contact-induced scaffold condensation, and that modulating IDR-IDR self-attraction is an effective strategy for achieving this fine tuning.

Multivalency of receptor binding provides a second control parameter. We systematically varied the PDZ valency of the hnRNPA1 IDR-containing scaffold by replacing one or more PDZ domains with a non-peptide binding mutant (see Methods).^41^ Progressively reducing the number of functional PDZ domains diminished contact-induced condensation, and the three-PDZ architecture was required to drive robust partner cell-induced condensation (**Fig. 2c, Extended Data Fig. 3b**). The expression levels of all PDZ scaffold variants are matched, ruling out the possibility that the observed condensation behaviors are caused by uncontrolled expression levels (**Extended Data Fig. 3d**).

We then asked whether the same design logic could be transferred to a different set of modules. For the receptor interaction, we replaced the extracellular GFP/αGFP cell-cell recognition interaction pairs with CD19 and an αCD19 single-chain antibody. For the intracellular scaffold-receptor interaction, we replaced the intracellular PDZ-peptide interaction with the orthogonal SYNZIP18-SYNZIP19 coiled-coil pair (**Fig. 2d**).^45^ Each SYNZIP scaffold protein was fused to a SNAP tag for visualization.^46^ A multivalent SYNZIP scaffold bearing DDX4 IDRs, which did not condense in the PDZ architecture, now underwent contact-induced enrichment with CD19-directed partner cells (**Fig. 2e**). The hnRNPA1 IDR-containing SYNZIP scaffold also showed induced condensation, whereas the FUS IDR-containing scaffold was strongly constitutive and the flexible-linker control did not condense (**Extended Data Fig. 4a, b**). The flow cytometry analysis shows that expression levels of the SYNZIP scaffold variants are matched to each other (**Extended Data Fig. 4c**). The change in DDX4 behavior between the PDZ and SYNZIP architectures suggests that cell contact-induced condensation is a context dependent property of the complete scaffold-receptor interaction network. Importantly, this context dependence is designable.

### Orthogonal synthetic scaffolds encode distinct synapses within a single cell

Natural cells often build compositionally different interfaces with different neighbors. Neurons, for example, can maintain molecularly distinct pre- and postsynaptic specializations at different contacts within the same cell.^3^ We therefore asked whether orthogonal scaffolds could allow one receiver cell to implement multiple partner-specific interfaces within the shared cytoplasm (**Fig. 3a**).

**Figure 3.**
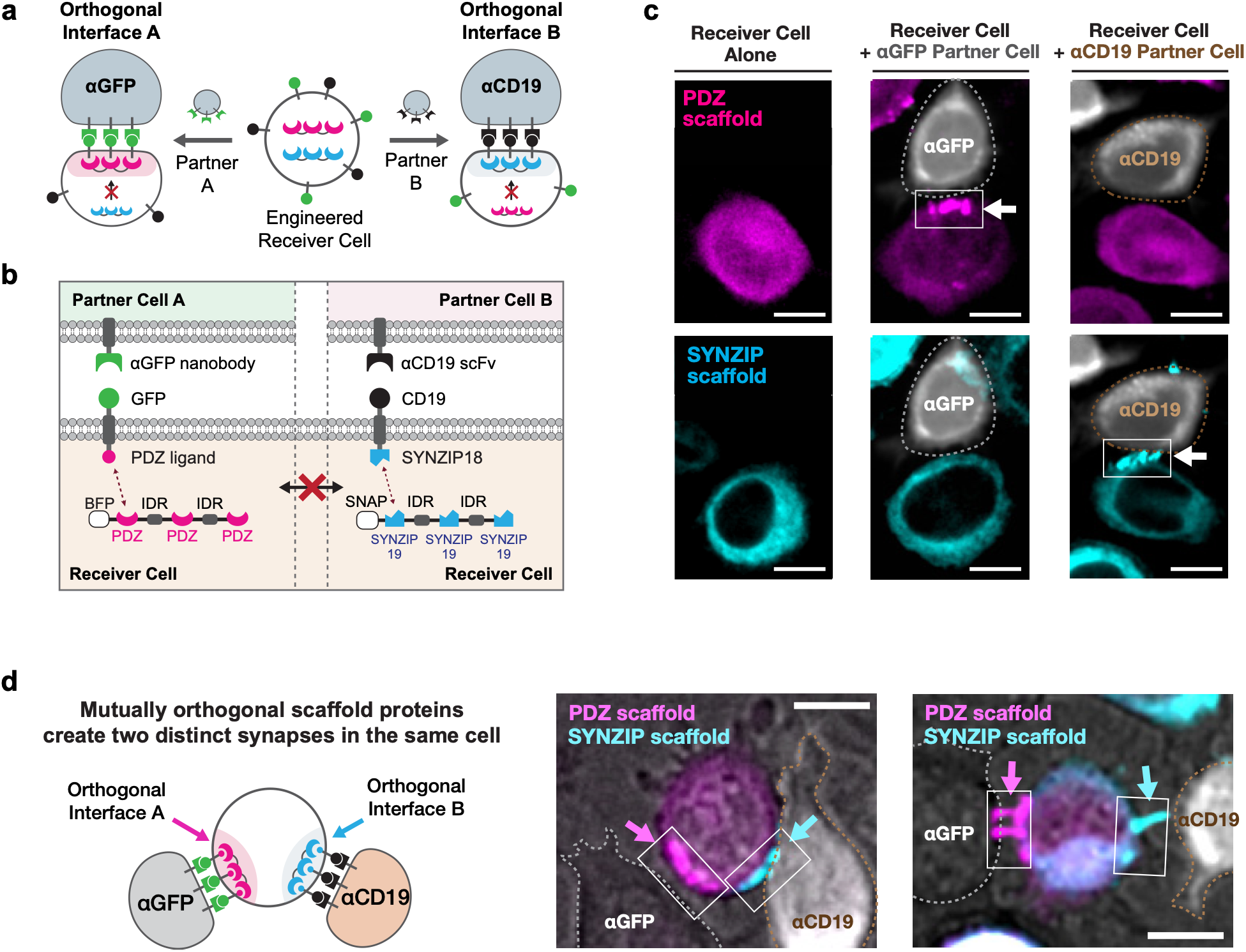
Mutually orthogonal scaffolds allow a single receiver cell to form distinct synapses with cognate partner cells. a, Schematic of an engineered receiver cell designed to assemble distinct synaptic complexes upon interaction with two different partner-cell types through orthogonal interfaces. b, Construct design of the two orthogonal synapses. The GFP/αGFP system recruits a PDZ scaffold, whereas the CD19/αCD19 system recruits a SYNZIP scaffold. c, Representative confocal images of dual-system receiver cells cultured alone or separately with each partner-cell type. PDZ scaffold is shown in magenta, SYNZIP scaffold in cyan and partner cells in white (labelled with CellTrace Far Red). Dashed outlines indicate partner-cell boundaries and arrows indicate scaffold enrichment. Scale bars, 10 µm; 2 h after co-culture. d, Representative confocal images of the receiver cell contacting both partner-cell types simultaneously, showing spatially distinct enrichment of the PDZ and SYNZIP scaffolds at their respective cognate interfaces. Scale bars, 10 µm. Quantification of interface specificity is shown in Supplementary Fig. 2.

We co-expressed the GFP/PDZ-hnRNPA1 IDR and CD19/SYNZIP-DDX4 IDR receptor/scaffold systems in the same receiver cell (**Fig. 3b**). Each synthetic receptor is linked to its corresponding intracellular scaffold-recognition module, and the two cytoplasmic scaffolds are designed to be mutually orthogonal. The dual-system receiver cells, when alone, show diffuse signals for both PDZ and SYNZIP scaffolds (**Fig. 3c**). When these receiver cells were co-cultured separately with αGFP or αCD19 partner cells for 2 hours, the cognate scaffold accumulated at the engaged interface while the other non-cognate scaffold remained diffuse (**Fig. 3c; Supplementary Fig. 2**). Thus, extracellular partner recognition was transmitted across the membrane into selective recruitment of a defined intracellular assembly.

The more stringent test was whether a single receiver could preserve selectivity while establishing contacts with both partner types simultaneously, the cellular topology akin to the pre- and post-synaptic densities in neurons. In mixed co-cultures of receiver cells with both αGFP and αCD19 partners, PDZ scaffold enrichment was restricted to αGFP interfaces, whereas SYNZIP scaffold enrichment was restricted to αCD19 interfaces (**Fig. 3d**). These results show that the modular nature of multivalent scaffold architecture can sufficiently reconstitute two distinct synaptic assemblies that coexist within one cytoplasm while maintaining their spatial segregation.

### Synthetic scaffold assemblies rewire junctional transport and orientation of cell division

We next tested whether cell contact-induced scaffold assemblies could functionally rewire cellular behaviors. Natural scaffolds organize channels, polarity complexes and signaling enzymes at defined cell-cell junctions, suggesting that a synthetic scaffold could also function as a local concentrator for effector proteins.^11,12^ We examined two qualitatively different outputs: intercellular molecular transfer through connexin channels and orientation of cell division through a recruited polarity factor.

For intercellular transfer, we used N2A neuroblastoma cells, which offer a gap junction-deficient background widely used to study exogenous connexin channels.^47^ We introduced a chimeric connexin-43 (Cx43) bearing HaloTag and a cognate PDZ-binding ligand (Cx43*; **Fig. 4a**). Here, connexins – which homo-oligomerize across cell-cell interfaces – act as the inducing receptor. We then asked whether the hnRNPA1 IDR-containing synthetic PDZ scaffold could assist the organization of Cx43* hemichannels at a cell–cell contact and facilitate the molecular transfer. Indeed, in cells expressing the hnRNPA1 IDR-containing PDZ scaffold, Cx43* and scaffold became co-enriched at interfaces.

**Figure 4.**
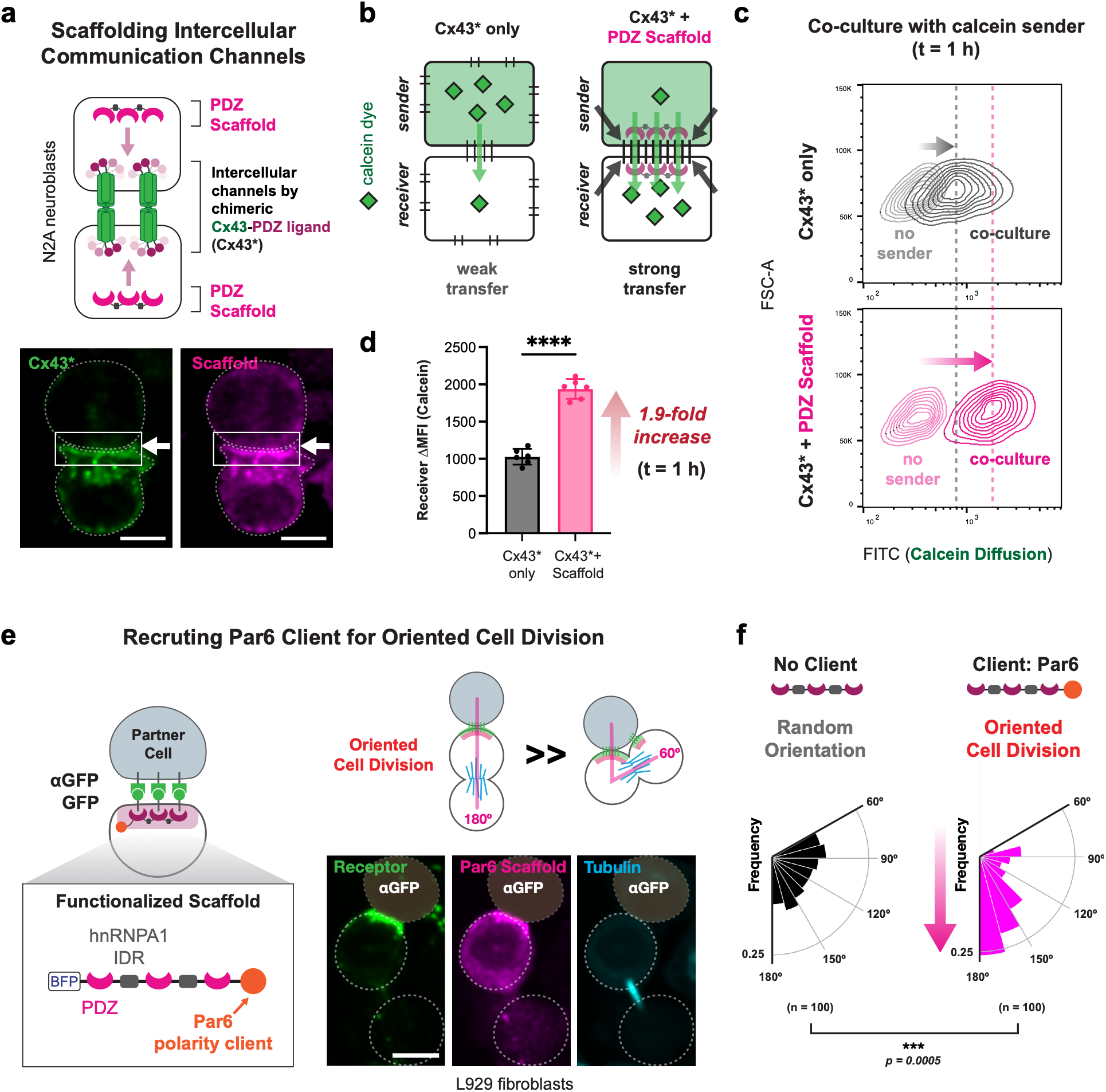
Engineered scaffold proteins rewire intercellular transport and orientation of cell division. a, Synthetic scaffolding of intercellular communication channels. In N2A neuroblastoma cells, a chimeric Cx43 protein bearing HaloTag and a PDZ-binding ligand (Cx43*) was co-expressed with the cognate hnRNPA1-containing PDZ scaffold. Representative confocal images show enrichment of Cx43* (green) and scaffold (magenta) at the cell–cell interface (fixed cells). Scale bars, 10 µm. b, Schematic of calcein dye transfer assay. Sender cells are loaded with calcein-AM and co-cultured with unlabeled receiver cells at 4:1 ratio. c, Representative receiver-cell flow-cytometry plots after 1 h of co-culture, comparing receiver cells expressing Cx43* alone with the receiver cells expressing Cx43* and the cognate PDZ scaffold. Cx43* expression matching is shown in Supplementary Fig. 2. d, Quantification of calcein dye transfer. The change in mean fluorescence intensity (MFI) of calcein in receiver cells were measured by flow cytometry analysis. Statistical significance was assessed by unpaired t-test (P < 0.0001). e, Schematic and confocal microscopy of a functionalized synapse designed to recruit the polarity protein Par6 and bias the orientation of cell division. Representative L929 cells show receptor (green), Par6-functionalized scaffold (magenta) and α-tubulin (cyan) at a partner-cell interface (fixed cells). Scale bar, 10 µm. f, Distribution of receiver-cell division orientation relative to the cell–cell interface with unfunctionalized or Par6-recruiting scaffold. n = 100 cells per condition. Statistical significance was assessed by Kruskal–Wallis (P = 0.0005).

We then measured their functional coupling with a calcein-transfer assay (**Fig. 4b**). Calcein-loaded sender cells were mixed with unlabeled receiver cells at 4:1 ratio, and dye accumulation in receivers was quantified by flow cytometry after one hour of co-culture. Cells expressing both Cx43* and the cognate PDZ scaffold accumulated more calcein than cells expressing Cx43* alone, with an approximately two-fold difference in receiver fluorescence (**Fig. 4c,d**). Cx43* expression was matched across the compared populations (**Supplementary Fig. 3**). These results demonstrate that the synthetic scaffold can colocalize with a junctional membrane protein Cx43* and facilitate an increase in molecular communication between cells.

We next tested whether a cytosolic signaling factor could be recruited as a client of the synthetic synapse. We focused on Par6, a component of the PAR polarity network that couples spatial cues to cytoskeletal and spindle organization.^48^ Recent work has shown that targeted enrichment of a single polarity factor Par6 is sufficient to decide the orientation of cell division.^49^ In our design, the synthetic scaffold was fused to Par6 while retaining its contact-induced assembly modules (**Fig. 4e**). Confocal microscopy shows that in engineered L929 fibroblast cells co-cultured with cognate partner cells, the receptor and Par6-functionalized scaffold accumulate at the synthetic interface.

The orientation of cell division was then quantified relative to the partner-cell interface. Receiver cells were sequentially treated with thymidine and nocodazole for cell cycle synchronization. The synchronized receiver cells were co-cultured with the αGFP partner cells, followed by nocodazole release and longitudinal microscopy imaging (see Method). Receiver cells bearing the unfunctionalized scaffold displayed a broad, random distribution of division angles, whereas recruitment of Par6 produced a significant bias in division orientation (n = 100 cells per condition; P = 0.0005 in the dataset shown; **Fig. 4f**). We therefore interpret the data as reorientation of the division axis by a localized cue triggered by Par6 enrichment. Together with the Cx43 experiment, these results show that a common scaffold architecture can be used to organize either a transmembrane channel or a cytoplasmic signaling module and thereby alter two distinct cell–cell functions that require localized enrichment of client proteins.

### Actin-regulatory clients sculpt synapse morphology

We next asked whether synthetic synapses could be functionalized to control the interface morphology by recruiting actin regulators. Actin-enriched protrusions and lamellar contacts are hallmarks of many natural synapses and are critical for processes such as immune surveillance, synaptic spine formation and remodeling, and epithelial junction formation.^2,50,51^ Filopodia and other narrow protrusions support exploratory contacts, whereas broad lamellar interfaces increase the area available for adhesion and signaling. Rho-family GTPases provide a compact way to control these morphologies: Cdc42 promotes filopodial actin organization and Rac1 promotes lamellipodial spreading.^52^ We therefore asked whether targeting Rho-family GTPase activators such as their guanine nucleotide exchange factors (GEFs) to the synthetic scaffold could couple partner recognition directly to local cytoskeletal remodeling.^53^

We fused the scaffold to catalytic domains from Intersectin (ITSN – a Cdc42 GEF) and TIAM1 (a Rac GEF). Intersectin can activate Cdc42 and promote N-WASP-dependent filopodial assembly, whereas TIAM1 is a Rac1 GEF that promotes membrane ruffling and lamellar protrusion.^54^ The receptor-binding PDZ domains and hnRNPA1 IDR architecture of the scaffold were otherwise retained. The partner cell expressed an αGFP receptor, but in this case we also appended an intracellular PDZ ligand and co-expressed the same GEF-functionalized PDZ scaffolds, in order to potentially drive condensation and GEF activity simultaneously on both sides of the synapse (**Fig. 5a**). Receiver cells were co-cultured with cognate partner cells for two hours, and receptor, scaffold and F-actin organization were examined at the resulting interfaces.

**Figure 5.**
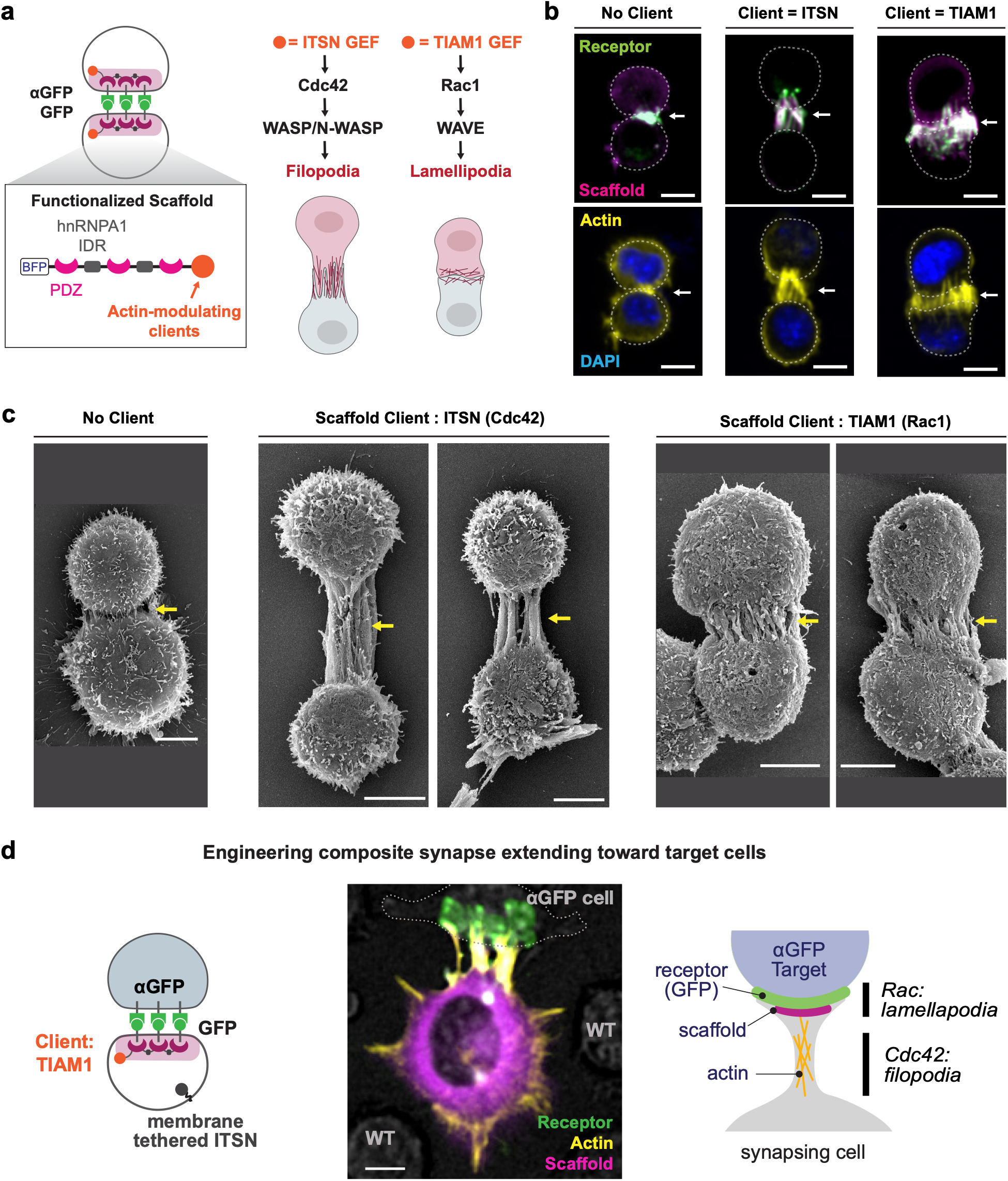
Recruitment of Rho-family GTPase GEFs to scaffold assembly remodels the morphology of engineered cell-cell synapses. a, Design of scaffolds functionalized with an ITSN GEF module to activate Cdc42 or a TIAM1 GEF module to activate Rac1. ITSN: Intersectin. Cells at both sides of cell-cell interaction are engineered with the GFP or αGFP receptor fused to PDZ ligand, together with functionalized PDZ scaffold. b, Representative confocal images of the cell-cell synapse formed between engineered L929 fibroblasts expressing the synthetic receptor and scaffold without a client, with scaffold-recruited ITSN or with scaffold-recruited TIAM1 (fixed cells). GFP Receptor is shown in green, scaffold in magenta, F-actin in yellow and nuclei in blue. Scale bars, 10 µm. 2 hours after co-culture. Quantification of cell–cell distance, interface width and actin enrichment is shown in Extended Data Fig. 5. c, Representative scanning electron microscopy images showing compact interfaces without a client, narrow protrusive interfaces with ITSN and broad lamellar interfaces with TIAM1. Scale bars, 5 µm. 2 hours after co-culture. d, Composite synapse in which membrane-tethered ITSN promotes exploratory protrusions and a TIAM1-functionalized synthetic synapse expands contacts selectively with cognate αGFP partner cells. Receptor is shown in green, F-actin in yellow and scaffold in magenta. Scale bar, 10 µm. 2 hours after co-culture.

The unfunctionalized scaffold (in both receiver and partner cell) generated a compact interface enriched in receptor and scaffold but with modest actin accumulation (**Fig. 5b**). Recruiting ITSN transformed this interface into distinct synaptic morphology with narrow, finger-like and actin-rich protrusions extending between the partner cells. Recruiting TIAM1 produced a very different synaptic morphology where contacts became broader and sheet-like, with actin enrichment distributed across an enlarged interface. Quantification confirmed significant changes in interface geometry and actin enrichment (**Extended Data Fig. 5**). Both ITSN and TIAM1 extended the lengths of cell-cell synapses quantified by the cell–cell distance metric (P < 0.0001 for each comparison). While ITSN made the interface narrower, TIAM1 enlarged the interface, consistent with the visually distinct narrow and broad contact morphologies (P = 0.0005 and 0.0043, respectively). Both ITSN and TIAM1 significantly enriched F-actin to the cell-cell synapses compared to the no-client control (P < 0.0001 for each comparison). Scanning electron microscopy independently revealed long, narrow protrusions in the ITSN condition and wide, flattened contacts in the TIAM1 condition (**Fig. 5c**). Because these structures were classified primarily by morphology, we refer to them as filopodia-like and lamellar rather than assigning a definitive ultrastructural identity.

We finally combined exploratory protrusion formation with partner-specific interface expansion to engineer the composite and targeted synapse structure, reminiscent of neuronal axon formation. A membrane-tethered ITSN module constitutively generated actin-rich filopodial projections from receiver cells. When these cells were engineered to also express the TIAM1-functionalized synthetic synapse, contacts with cognate αGFP partner cells developed into extended actin-rich interfaces, whereas neighboring cells lacking the cognate ligand (WT L929) did not elicit synapse formation (**Fig. 5d**). This composite synapse illustrates how distinct cytoskeletal modules can be layered onto the scaffold-driven morphology: one module (Cdc42 mediated) increases the opportunity to drive encounters between neighboring cells, whereas the synapse formation module (Rac mediated) selectively stabilizes and expands the morphological size of contacts between partner cells.

## Discussion

Here, we establish a physical mechanism for organizing *de novo* cell-specific cell-cell synapses, providing a reductionist perspective on how native junctions assemble. Central to this mechanism is the modularity of multivalent scaffold proteins. Our simulations reveal that local enrichment of cognate receptors can drive the formation of membrane-localized scaffold condensates. However, scaffold condensation exhibits ultra sensitivity to receptor density only when scaffold self-association is maintained within an optimally tuned regime. We show that varying the IDRs and binding valency of a minimal scaffold provides a pair of design handles that enable the achievement of the fine-tuning requirement. By modulating these parameters within an architecture comprising three PDZ domains and two IDRs, we successfully engineered synthetic synapses that undergo switch-like assembly strictly upon partner cell contact. This synthetic system reconstitutes several fundamental features of natural junctions: contact restriction, cooperative enrichment, partner selectivity, and modular client recruitment.

The modularity of the synthetic synapse architecture also enables the rational construction of orthogonal cell–cell interfaces and reveals how heterotypic receptor-scaffold binding affinity shapes condensate behavior. We generated functional synapses operating across vastly different affinity regimes: a PDZ system driven by weak, micromolar binding (Kd ∼7.3 µM)^41^ and a SYNZIP system driven by strong, nanomolar binding (Kd < 10 nM).^45^ The ability of the same three-domain, two-IDR architecture to function across nearly three orders of magnitude in binding affinity underscores the robustness of scaffold-mediated condensation. Moreover, strengthening heterotypic receptor–scaffold binding in the SYNZIP system expands the reach of weakly condensing IDRs, allowing them to undergo contact-induced assembly as was shown for DDX4 IDRs and the flexible GS linkers. These IDRs, which form mostly diffuse distributions, can be coaxed to undergo meaningful, context-dependent degrees of contact-induced condensates (66% and 24% of total cell populations, respectively, **Fig. 2c, Extended Data Fig. 3c**, and **Extended Data Fig. 4b**). Thus, the strength of heterotypic receptor–scaffold binding plays a critical role in driving interface formation and tuning its physical properties and mechanisms, which remain an active area of investigation for future studies.

A key advantage of the modular architecture is spatial multiplexing. By using orthogonal receptor–scaffold pairs, one cell can maintain different molecular assemblies at different interfaces even though the scaffold proteins share the same cytoplasm. This creates a form of localized and polarized partner cell-specific response at the protein-assembly level. This is conceptually appealing because the topology is reminiscent of how neurons create pre- and post-synaptic densities within one cell depending on the interacting partner neurons. At the engineering level, this distinguishes our platform from other synthetic receptor systems that primarily route contact information into transcriptional programs or whole cell state changes.^55–58^ Here, the immediate output is a supramolecular structure that remains physically associated with the site of engagement. Such spatial selectivity provides a basis for achieving functional outputs that depend on subcellular localization of signaling molecules, such as membrane channels, polarity regulators or the cytoskeleton.

Our functional experiments further show that the multivalent scaffold protein not only provides the general physical organizing principle but also serves as the minimally sufficient system for functional synapse formation. Concentrating chimeric connexin-43 increased net dye transfer, recruiting Par6 biased the orientation of division, and recruiting Rho-family GEFs generated distinct interfacial morphologies. These outputs span membrane transport, cell polarity and actin mechanics, suggesting that many additional effectors could in principle be localized using the same core architecture, similarly as natural cell-cell synapses carry out diverse physiological functions to mediate complex multicellular communication.

Finally, because recognition modules, scaffold interactions and functional clients can be exchanged independently, the modular architecture provides a compact route to engineering cell–cell interfaces with programmable composition and behavior. This finding is consistent with the evolution of condensate forming scaffold proteins in early metazoans, likely as a means to create specific cell-cell interfaces, far before there were specialized cells such as neurons. In the future, these engineered cell-cell synapses could be applied to address therapeutic problems such as precise control of the interactions of immune cells and neural cells, or to potentially allow the design of custom multi-cell circuits and networks.^59^

## Methods

### Materials

L929 mouse fibroblasts (CCL-1) were obtained from the American Type Culture Collection (ATCC). Neuro-2a (N2A) cells were provided by the laboratory of Orion Weiner. Lenti-X 293T cells (Takara Bio) were used for lentiviral production. Cells were cultured in Dulbecco’s modified Eagle medium containing GlutaMAX (Gibco) supplemented with 10% fetal bovine serum (Gibco). TrypLE Express and Opti-MEM (Gibco) were used for cell dissociation and transfection, respectively. TransPlus Virus Transduction Enhancer (Alstem) was used during lentiviral transduction. Poly-D-lysine (Sigma Aldrich) was used to coat culture surfaces when indicated. Assays in 384-well format were performed in Corning BioCoat Poly-D-Lysine 384-well Black/Clear Flat Bottom TC-treated Microplates.

Oligonucleotides were purchased from Integrated DNA Technologies, and synthetic DNA fragments were purchased from Twist Bioscience. Q5 High-Fidelity 2X Master Mix (New England Biolabs) was used for PCR amplification. DNA fragments were assembled using In-Fusion Snap Assembly Master Mix (Takara Bio) or Golden Gate assembly with either the BsmBI-v2 Golden Gate Assembly Kit or Esp3I, T4 DNA Ligase Buffer, and CutSmart Buffer (NEB), as appropriate. Stellar chemically competent *Escherichia coli* (Takara Bio) were used for plasmid propagation. Plasmid DNA was purified using QIAprep miniprep kits, and DNA fragments were purified using QIAquick gel extraction kits and spin columns (Qiagen).

Lentiviral particles were produced using the packaging plasmids pMD2.G and p8.91 and TransIT-VirusGEN Transfection Reagent (Mirus Bio). Viral supernatants were filtered through 0.45-µm filters and concentrated using Lenti-X Concentrator (Takara Bio).

Partner cells were distinguished using CellTrace Yellow or CellTrace Far Red Cell Proliferation Kits (Thermo Fisher Scientific). CellTrace Calcein Red-Orange AM (Thermo Fisher Scientific) was used in Connexin experiments. SNAP-tagged proteins were labeled with SPY555-BG (Spirochrome), and filamentous actin was labeled with SPY650-FastAct (Spirochrome). Live cell microtubule was labeled with SPY650-tubulin (Spirochrome). Verapamil (Sigma-Aldrich) was included during labeling where indicated. Surface expression was measured using an Alexa Fluor 647 DYKDDDDK Epitope Tag Antibody (1042E) (R&D Systems) or a PE-conjugated Myc-Tag (9B11) Mouse Monoclonal Antibody (Cell Signaling Technology). For fixed samples, DAPI (Thermo Fisher) was used for nuclear staining and ActinRed 555 ReadyProbes Reagent (Thermo Fisher) was used for actin staining following the recommended protocol. For calcein dye transfer, calcein-AM (eBioscience) was used. Thymidine (Sigma Aldrich) and nocodazole (Sigma Aldrich) were used for cell cycle arrest protocols.

### Equipment

Cell sorting was performed using the FACSAria II Cell Sorter or Sony SH800S Cell Sorter. Flow cytometry was performed with the BD LSRFortessa Cell Analyzer or the BD FACSymphony A1 Cell Analyzer. Confocal microscopy was performed using the Opera Phenix automated spinning-disk confocal microscope with a ×40 water-immersion objective in 384-well plates. Structured illumination microscopy was performed with the DeltaVision OMX SR Super-resolution Microscope. Scanning electron microscopy was performed with a Phenom Pharos G2 Desktop FEG-SEM.

### Construct design and cloning

Coding sequences for naturally occurring proteins were obtained from UniProt (https://www.uniprot.org)^60^ and relevant sequences are provided in Supplementary Table 2. Synthetic protein sequences were derived from the following publications: anti-GFP nanobody,^40^ PDZ interaction modules (Peptide 24 and Erbin PDZ 24 for the PDZ scaffolds, and Erbin PDZ 12 for the non-binding mutant),^41^ SYNZIP interaction modules (SYNZIP18, 19),^45^ SNAP tag,^46^ HaloTag,^61^ and Fibcon.^62^ Unless otherwise noted, coding sequences were codon-optimized for expression in mammalian cells using Integrated DNA Technologies’s codon optimization tool.

All constructs contained a Kozak consensus sequence and were cloned into a pHR lentiviral transfer vector. Expression was driven by either the phosphoglycerate kinase (PGK) promoter or the spleen focus-forming virus (SFFV) promoter, as specified in Supplementary Table 1. Surface-receptor constructs contained an IRES2 element, and scaffold constructs contained a nuclear export signal to reduce nuclear localization. Plasmids were assembled using In-Fusion cloning or Golden Gate assembly and propagated in Stbl3 chemically competent E. coli (Macro lab). Final plasmid sequences were verified by whole plasmid sequencing by Angstrom Innovation, Plasmidsaurus, or Genewiz. Complete plasmid architectures and sequences are provided in Supplementary Table 1.

### Lentiviral production

Lenti-X 293T cells were seeded in six-well plates at 7.0 × 10^5^ cells per well and allowed to attach overnight. The following day, cells were co-transfected with the transfer plasmid of interest and the packaging plasmids pMD2.G and p8.91 using TransIT-VirusGEN Transfection Reagent according to the manufacturer’s instructions. Viral supernatants were collected 48 h after transfection, passed through a 0.45-µm filter, and concentrated tenfold using Lenti-X Concentrator according to the manufacturer’s instructions.

### Cell culture and generation of engineered cell lines

L929, N2A, and Lenti-X 293T cells were maintained in DMEM GlutaMAX supplemented with 10% FBS. To generate engineered cell populations, 1.0 × 10^5^ cells were seeded per well in 12-well plates and transduced with lentiviral supernatant in the presence of TransPlus Virus Transduction Enhancer according to the manufacturer’s protocol. After 24 h, the viral medium was replaced with fresh growth medium. Cells were analyzed or sorted 48 hours after transduction.

Cells were sorted based on either a fluorescent protein encoded by the construct or staining of the appropriate extracellular epitope tag. The antibodies used for sorting are Alexa Fluor 647 DYKDDDDK Epitope Tag Antibody (1042E) (R&D Systems) or a PE-conjugated Myc-Tag (9B11) Mouse Monoclonal Antibody (Cell Signaling Technology) at a 1:200 dilution. To generate expression-matched populations for the IDR and valency experiments, cells were transduced with multiple viral titers. Expression distributions were measured by flow cytometry, and the populations with the most closely matched median fluorescence intensity were selected for subsequent assays.

### Cell contact assay

Receiver cells were seeded in poly-D-lysine-coated, black, clear-bottom 384-well plates at 4.0 × 10^3^ cells per well and allowed to settle for 1 h. Partner cells were then added at 4.0 × 10^3^ cells per well, producing a 1:1 partner-to-receiver ratio. Co-cultures were incubated for 2 h at 37 °C before imaging.

### Live fluorescent labeling and confocal imaging

SNAP-tagged scaffold proteins were labeled with SPY555-BG according to the manufacturer’s instructions. Filamentous actin was labeled with SPY650-FastAct according to the manufacturer’s instructions. Microtubule was labeled with SPY650-tubulin according to the manufacturer’s instructions. Verapamil was included during labeling according to the dye manufacturer’s recommended protocol to reduce dye efflux. Partner cells were labeled with CellTrace Yellow or CellTrace Far Red according to the manufacturer’s instructions. Confocal images were acquired in 384-well plates using an Opera Phenix system (Revvity) using a 40× water-immersion objective. All confocal microscopy images are from live cell imaging unless noted otherwise.

### Classification of condensation regimes

A condensate was operationally defined as a discrete area of scaffold enrichment with a fluorescence intensity at least 1.5-fold greater than the remaining cellular scaffold signal. Receiver cells were assigned to one of three mutually exclusive phenotypic classes: no condensation, interface-restricted condensation, or constitutive condensation. Cells were classified as exhibiting no condensation if no condensate is observed with and without co-culturing with the cognate partner cells. Cells were labeled as exhibiting interface-restricted condensation when no condensate is observed by themselves, but at least one qualifying condensate was detected at a partner-receiver interface after co-cultured with the cognate partner cells. Cells were classified as exhibiting constitutive condensation when at least one qualifying condensate was present before being co-cultured with the partner cells. At least 50 cell-cell contacts were scored per condition in each experiment. A contact was defined by the physical proximity of less than 5 µm.

### Immunofluorescence staining

Cells were fixed in 4% paraformaldehyde in phosphate-buffered saline (PBS) for 15 min at room temperature and washed three times with PBS. For intracellular targets, cells were permeabilized for 1 h at room temperature in PBST (0.25% Triton X-100 in PBS). Samples were then incubated in blocking buffer (0.1% BSA, 0.25% Triton X-100 in PBS) for 3 h at room temperature or overnight at 4 °C with gentle rocking. Fluorophore-conjugated antibodies, were diluted in blocking solution and incubated with samples overnight at 4 °C with gentle rocking. Samples were then washed three times in PBST, with a 15-min incubation during each wash. Secondary-stained samples were washed three additional times before imaging. The following antibodies and dilution were used: anti-Connexin 43 rabbit monoclonal antibody 3O6 CoraLite Plus 488 (Thermo Fisher, 200x), V5 Tag Monoclonal Antibody TCM5 PE eBioscience (Invitrogen, 200x) V5 Tag Monoclonal Antibody Alexa Fluor 647 (Invitrogen, 200x), and Alexa Fluor 647 Anti-alpha Tubulin antibody EP1332Y (abcam, 500x).

### Scanning electron microscopy

Round glass coverslips were placed in 24-well plates and coated with 100 µg/mL poly-D-lysine for at least 2 h at 37 °C or overnight at 4 °C. Receiver cells and partner cells were each seeded at 1.0 × 10^5^ cells per coverslip, for a total of 2.0 × 10^5^ cells per well, and cultured for 1 hour before fixation. Samples were fixed in 1.5% paraformaldehyde for 30 min.

Fixed samples were postfixed in 1% osmium tetroxide in water for 1 h at room temperature while protected from light. Samples were dehydrated through a graded ethanol series consisting of 25%, 50%, 75%, and 100% ethanol, with a 15-min incubation at each step. Samples were dried by critical-point drying with liquid CO2 using a Leica EM CPD300 critical-point dryer. Dried samples were sputter-coated with iridium using a Leica EM ACE600 Sputter Coater. Images were acquired using a Phenom Pharos G2 Desktop FEG-SEM.

### Calcein dye transfer assay

The functional assessment of junctional intercellular communication was adapted from previously established preloading dye transfer assays.^63^ Sender and receiver N2A cell populations were independently seeded at a density of 1 × 10⁶ cells per well in 6-well plates and incubated at 37 °C for 15 h. A staining solution was prepared in PBS containing 2.5 µM Calcein-AM and 1.25 µM CellTrace Far Red. The medium of the sender cells was replaced with 1 mL of the staining solution, followed by a 20-min incubation at 37 °C. Subsequently, 2 mL of complete medium was added, and cells were incubated for an additional 20 min at 37 °C. To remove residual extracellular dye, the sender cells were washed three times with PBS, replenished with complete medium, and incubated for 30 min. Both sender and receiver populations were then dissociated using Trypsin-EDTA, centrifuged, and resuspended in complete medium. Co-cultures were prepared by plating 8 × 10⁴ sender cells and 2 × 10⁴ receiver cells per well in a 96-well plate. After a 1-h incubation at 37 °C, the plates were centrifuged at 200 × g for 5 min. The medium was gently aspirated, and the cells were trypsinized and resuspended for immediate flow cytometry analysis.

### Quantification of cell division orientation

Receiver L929 cells were synchronized using a sequential thymidine-nocodazole block and mitotic shake-off protocol, adapted from a previously described protocol.^64^ Briefly, 4 × 10⁵ receiver cells were seeded into a 6-well plate and treated with 2 mM thymidine in complete medium for 10 h at 37 °C. The thymidine-containing medium was aspirated, and the cells were washed and allowed to recover in fresh complete medium for 3 h at 37 °C. Cells were subsequently treated with 100 ng/mL nocodazole for 6 h at 37 °C to induce prometaphase arrest. To isolate the synchronized mitotic population, the medium containing dead cells was gently aspirated and replaced with fresh nocodazole-containing medium (100 ng/mL), followed by mechanical detachment via mitotic shake-off. The harvested cell suspension was labeled with SPY650-tubulin supplemented with 10 µM verapamil. Labeled receiver cells were then seeded at a density of 3 × 10³ cells per well in a 384-well plate and incubated for 30 min at 37 °C. Partner cells, previously labeled with CellTrace Far Red according to the manufacturer’s instructions, were added to the receiver cells at a density of 3 × 10³ cells per well and co-cultured for 1 h at 37 °C. The nocodazole arrest was released by washing the co-cultures three times with complete medium, after which longitudinal confocal microscopy imaging was initiated at 1-h intervals. Mitotic progression typically resumed immediately upon release, with anaphase and cytokinesis observed at the 3–4 h time point. The division orientation angle was quantified from these confocal images by determining the axis of the mitotic spindle (visualized by SPY650-tubulin) relative to the cell–cell interface (indicated by the GFP-fused receptor enrichment).

### Coarse-grained molecular dynamics simulations

#### Model Description

We performed coarse-grained (CG) molecular dynamics (MD) simulations using the LAMMPS (<u>L</u>arge-scale <u>A</u>tomic/<u>M</u>olecular <u>M</u>assively <u>P</u>arallel <u>S</u>imulator)^33^ to model the cooperative condensation of scaffold proteins at a receptor-decorated membrane surface. Each scaffold molecule is represented as a linear homopolymer chain of 25 beads (atom type 1) connected by finitely extensible nonlinear elastic (FENE)^65^ bonds. Our choice for the number of beads was made to mimic the number of cohesive motifs (stickers) in PLCDs. Surface receptors are represented as individual beads (atom type 2) that diffuse laterally within a quasi-two-dimensional slab at the upper boundary of the simulation box, representative of receptors on a cell membrane.

The simulation box has dimensions 60σ × 60σ × 70σ with periodic boundary conditions in x and y and fixed (non-periodic) in z. Here, σ is the bead diameter. Scaffold chains are confined between repulsive walls at z = −32σ and z = +32σ, giving an effective scaffold volume of 60σ × 60σ × 64σ. Receptors are confined to a slab between z = 32σ and z = 35σ by separate walls. The membrane surface area is (60σ)^2^ = 3600σ^2^, and receptor density is defined as D_R_ = N/3600 molecules/σ^2^, where N is the number of receptors. A receptor density of D_R_ = 1 molecule/σ^2^ corresponds to 3600 receptors on the surface. The system contains 200 scaffold chains (5000 scaffold beads).

#### Interaction potentials used

Non-bonded interactions between all bead pairs are modeled using the Lennard-Jones (LJ) potential^66^ with a shifted cutoff,

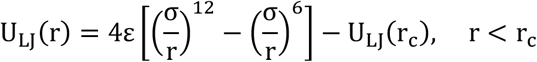

where σ = 1.0 is the bead diameter and the potential is shifted to zero at the cutoff distance r_c_. Three pairwise interactions are defined: the tunable scaffold-scaffold homotypic attraction (ε_S/S_) with a cutoff r_c_ = 2.5σ, the receptor-scaffold heterotypic attraction (ε_R/S_) fixed at 15 kT, and with a longer cutoff r_c_ = 4.0σ, and a pure Weeks-Chandler-Andersen (WCA) repulsion between the receptor beads to prevent their overlap, with ε_R/R_ = 1.0 kT and r_c_ = 2(1/6)σ ≍ 1.1225σ.

Bonds between consecutive beads within a scaffold chain are modeled using the FENE potential with parameters k = 10.0, r_0_ = 2.0σ, and a built-in LJ repulsion with ε = 1.0 kT and σ = 1.0. The maximum bond extension is r_0_ = 2.0σ and the equilibrium bond length is approximately 1.0σ.

Scaffold molecules (chains) are confined by repulsive LJ walls (ε = 1.0 kT, σ = 1.0, r_c_ = 2(1/6)σ) at z = ±32σ. Receptors are confined by similar walls at z = 32σ and z = 35σ. All pair potentials are shifted to zero at their respective cutoffs.

#### Simulation protocol

An initial configuration of 200 scaffold chains (5000 beads) was setup in a receptor-free simulation box (N = 0) with all pair interactions set to WCA (purely repulsive, ε = 1.0 kT, r_c_ = 2(1/6)σ) to generate a well-dispersed, non-aggregated starting state. The energy of the system is minimized by iteratively adjusting atomic positions using the conjugate gradient algorithm until the potential energy reaches a local minimum, for a maximum of 100 iterations (steps) or 1000 force evaluations, stopping sooner if the energy or force tolerances are met. The equilibration consisted of 10,000 steps of velocity-limited NVE integration followed by 1,000,000 steps of NVE + Langevin dynamics at kT = 1.0 with a damping parameter of 1.0 τ. The resulting equilibrated configuration served as the common starting point (using a LAMMPS restart file) for all the production simulations.

For each production simulation, receptors were randomly placed in the membrane slab (z = 32.2σ to z = 34.8σ) using the LAMMPS create_atoms command with a fixed random seed for receptor placement. Production runs began with 10,000 steps of velocity-limited NVE integration to relax any initial overlaps introduced by receptor placement, followed by NVE + Langevin dynamics at kT = 1.0 with a damping parameter of 100.0 τ and a timestep of Δt = 0.005 τ. Each production run involves 5×10^7^ time steps (250,000 τ).

Convergence of MD simulations was verified by monitoring the average potential energy (PE) per bead and the fraction of scaffold in the largest connected cluster (f_lc_) as functions of simulation time (see **Supplementary Fig. 1**). For each combination of ε_S/S_ and D_R_ (or equivalently N), five independent replicate simulations were performed. Replicates differ only in the random seed that sets the initial velocities. Using the randomized initial velocities and the Langevin thermostat, leads to independent trajectories, albeit from the same initial configuration. Quantities reported in the analyses were averaged over the final 10 frames of the fully converged simulations for each replicate (50 measurements per condition), and error bars represent the standard deviation across all 50 measurements. The simulation parameters are summarized in Supplementary Table 2.

#### Analysis: (a) Fraction of scaffold in largest cluster (f_lc_)

Scaffold beads (type 1 only; receptors excluded) were clustered using a distance-based criterion with a cutoff of 2.0σ, matching the FENE bond maximum extension r_0_. Periodic boundary conditions were applied in x and y (but not z) when computing inter-bead distances. Connected components were identified from the resulting neighbor graph using sparse matrix methods (scipy.sparse.csgraph.connected_components). The largest cluster fraction f_lc_ is the number of beads in the largest connected component divided by the total number of scaffold beads (5,000).

#### (b) Threshold value of ε_S/S_ for bulk condensation

We determined the threshold value for ε_S/S_ beyond which by performing a sweep of ε_S/S_ from 0.1 to 1.0 kT at minimal receptor density (N = 1, D_R_ = 0.0003 molecules/σ^2^). The order parameter f_lc_ was computed at each ε_S/S_ and fitted with a sigmoidal function: f_lc_(ε_S/S_) = f_lc,dispersed_ + [(f_lc,condensed_ − f_lc,dispersed_) / (1 + exp(−k (ε_S/S_ − ε_S/S,critical_)))], where f_lc, dispersed_ and f_lc,condensed_ are the plateau values in the dispersed and fully condensed limits, and k is the steepness of the sigmoidal transition. The fitted threshold value is ε_S/S,critical_ ≍ 0.4873 kT. Below this value, scaffold remains dispersed and above it, constitutive condensation occurs regardless of receptor density.

#### Visualization

For visualization in OVITO,^67^ LAMMPS data files and trajectories were post-processed to assign scaffold beads in the largest cluster to a distinct particle type (type 3), yielding three visually distinguishable species: scaffold not in the largest cluster (type 1), receptors (type 2), and scaffold in the largest cluster (type 3).

### Bioinformatics analysis of the domain architectures in cell-cell junction proteins housing long IDRs

IDR containing proteins were extracted using the method described in earlier work.^68^ Briefly, MobiDB^69,70^ was used to predict all IDRs of length ≥ 30. Then a filtering was performed to extract IDRs of length ≥ 70 and non-linker IDRs of length ≥ 50. We refer to these IDRs as “long IDRs”. This criterion was chosen to select IDRs that are likely to have functional roles within the context of the full protein. The specific length cutoff for the number of residues is similar to the length of folded domains. Next, we extracted domain information for each protein in the human proteome using UniProt.^60^ If the length of the full protein in the IDR set did not match the length of the full protein in the domain set, due to a change in the main UniProt isoform between data extraction, the given protein was also filtered out. This avoids any potential domain / IDR mismatches based on the specific isoform used. Overall, we find 6269 unique proteins housing long IDRs (IDRs ≥ 70 and non-linker IDRs ≥ 50).

The list of proteins known to localize to cell junctions was extracted from UniProt using the GO term 0030054 for cell junctions.^71,72^ This list of proteins was further filtered to remove any proteins that had a transmembrane domain using the keyword for transmembrane (KW-0812) in UniProt. Overall, we find 1510 unique cell junction proteins without transmembrane domains. Additionally, 625 unique cell junction proteins without transmembrane domains also house long IDRs.

Using the extracted UniProt domain information we extracted 189 unique domains in the 625 unique cell junction proteins housing long IDRs. We then counted the number of unique proteins housing at least one copy of each unique domain. Of the 625 proteins, 52 house at least one PDZ domain, with proteins housing PDZ domains being the third most common, following only proteins housing SH3 and PH domains (**Extended Data Fig. 2b**).

Lastly, we analyzed the domain architecture of the 52 PDZ containing proteins. Specifically, we counted the number of protein binding domains and long IDRs in each protein. To determine which domains were protein binding domains we first extracted all domains within the 52 PDZ containing proteins. The protein binding domains were assigned based on Uniprot domain features. We found that cell junction proteins housing long IDRs and at least one PDZ domain have a mode of three binding domains and two long IDRs.

## Data Availability

Experimental data supporting the conclusions of this study are available within the Article and the Supplementary Information. All databases used in this study are publicly available. For identifying protein sequences and domain architecture, the Universal Protein Resource (https://www.uniprot.org/) was used. MobiDB was used to identify predicted IDRs,^69,70^ and GO was used for identifying subcellular localization.^71,72^ Source data are provided with this paper.

## Code Availability

Exemplary simulation code is provided at [https://github.com/Pappulab/Synthetic_Synapse].

## Acknowledgements

We thank H. Scott, K. H. Kim and the members of the Lim laboratory, Cell Design Institute, and members of the Pappu laboratory for discussions, assistance and advice. We thank D. Mullins for helpful discussions and sharing instruments for electron microscopy. N.Y.S. is a Damon Runyon Quantitative Biology Fellow supported by the Damon Runyon Cancer Research Foundation (DRQ, 16-23). This work was supported by grants from the US National Institutes of Health (R01NS121114 to RVP, U54CA244438 and U01CA265697 to WAL), the US National Science Foundation (MCB-2227268 to RVP, DBI-1548297 to WAL), the St. Jude Research Collaborative on the Biology and Biophysics of RNP Granules (RVP), and the Mark Foundation for Cancer Research (RVP).

## Author Information

Gaurav Chauhan

Present address: Department of Chemical Engineering, Indian Institute of Technology, Indore, India

## Contributions

NYS, RVP, and WAL designed research. NYS and AMG cloned plasmids, generated cell lines, performed experiments, and analyzed data. GM and GC designed and performed molecular dynamics simulations. KMR performed bioinformatics analysis. All authors worked together to analyze the data. NYS, RVP, and WAL wrote the paper with inputs from other authors.

## Supplementary Information

**Supplementary Figure 1.** Illustration of the convergence of the MD simulations (linked to main Figure 1c-f and Extended Figure 1b).

**Supplementary Figure 2.** Partner cells expressing different extracellular ligands induce condensation of the specific corrsponding scaffold proteins.

**Supplementary Figure 3.** Flow cytometry analysis of chimeric connexin 43 (Cx43*) expression levels in N2A neuroblast cells following cell sorting and Halotag labeling.

**Supplementary Table 1.** List of plasmids and features

**Supplementary Table 2.** Simulation parameters

**Supplementary Movie 1 (linked to Extended Data Fig. 1b).** Molecular dynamics simulation of strong homotypic scaffold (εS/S = 3.0 kT) and low receptor density (DR = 0.14).

**Supplementary Movie 2 (linked to Extended Data Fig. 1b).** Molecular dynamics simulation of strong homotypic scaffold (εS/S = 3.0 kT) and high receptor density (DR = 0.83).

**Supplementary Movie 3 (linked to Extended Data Fig. 1b).** Molecular dynamics simulation of intermediate homotypic scaffold (εS/S = 0.45 kT) and low receptor density (DR = 0.14).

**Supplementary Movie 4 (linked to Extended Data Fig. 1b).** Molecular dynamics simulation of intermediate homotypic scaffold (εS/S = 0.45 kT) and high receptor density (DR = 0.83).

**Supplementary Movie 5 (linked to Extended Data Fig. 1b).** Molecular dynamics simulation of weak homotypic scaffold (εS/S = 0.1 kT) and low receptor density (DR = 0.14).

**Supplementary Movie 6 (linked to Extended Data Fig. 1b).** Molecular dynamics simulation of weak homotypic scaffold (εS/S = 0.1 kT) and high receptor density (DR = 0.83).

## Source Data

Source Data Fig. 1

Source Data Fig. 2

Source Data Fig. 4

Source Data Extended Data Fig. 1

Source Data Extended Data Fig. 2

Source Data Extended Data Fig. 3

Source Data Extended Data Fig. 4

Source Data Extended Data Fig. 5

## Extended Data Figures and Tables

**Extended Data Figure 1.**
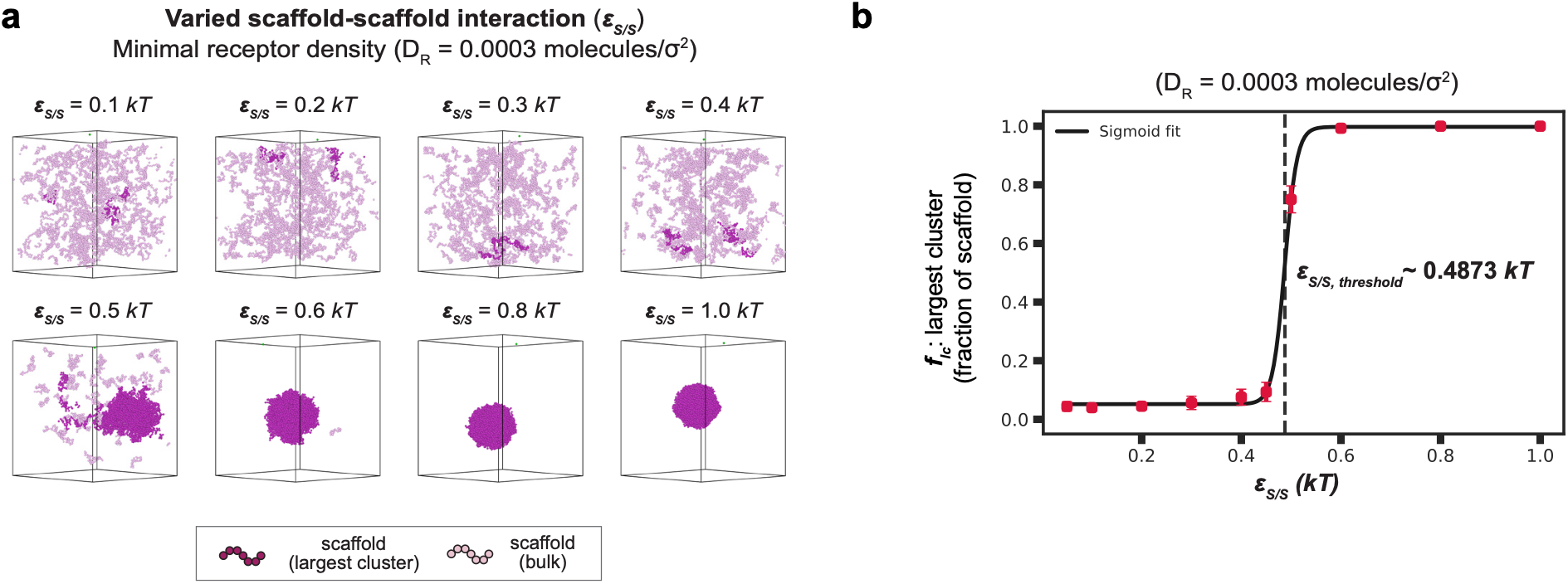
Simulation results of scaffold condensation (linked to main Fig. 1c-f). a, Representative simulation snapshots of the final frame showing scaffold condensation with varied scaffold-scaffold self-interaction strengths with minimal receptor densities. Here, ε_S/S_: homotypic scaffold-scaffold interaction and D_R_: the receptor density is kept at 0.0003 molecules/σ^2^, where σ: the effective diameter of each receptor. b, Determination of the critical ε_S/S_ where bulk condensation of scaffold is observed with minimal receptor presentation density (D_R_ = 0.0003 molecules/σ^2^), using a sigmoidal fitting function. This value is ε_S/S_,_threshold_ = 0.4873 kT, and above this threshold value, constitutive condensation is always observed. Here, f_lc_: fraction of scaffold molecules in the largest connected cluster.

**Extended Data Figure 2.**
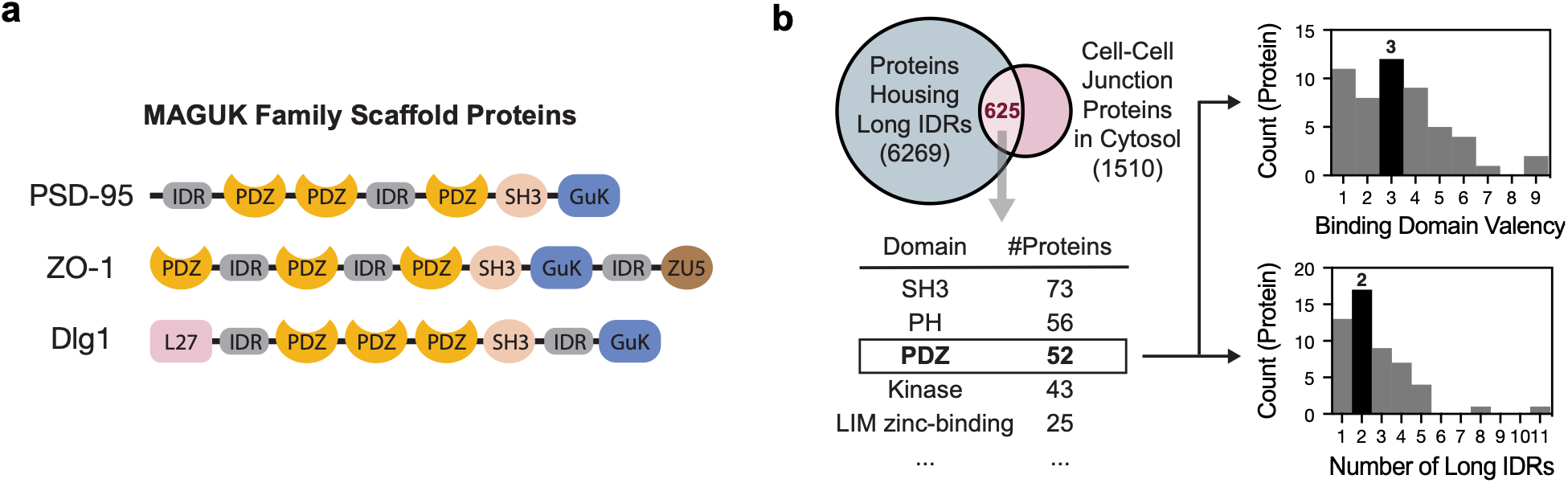
Basis of minimal scaffold design (linked to main Fig. 2a). a, Examples of MAGUK family scaffold proteins characterized by multiple protein interaction domains and intrinsically disordered regions (IDRs). b, Bioinformatics analysis of the domain architectures present in cell-cell junction proteins housing long IDRs.

**Extended Data Figure 3.**
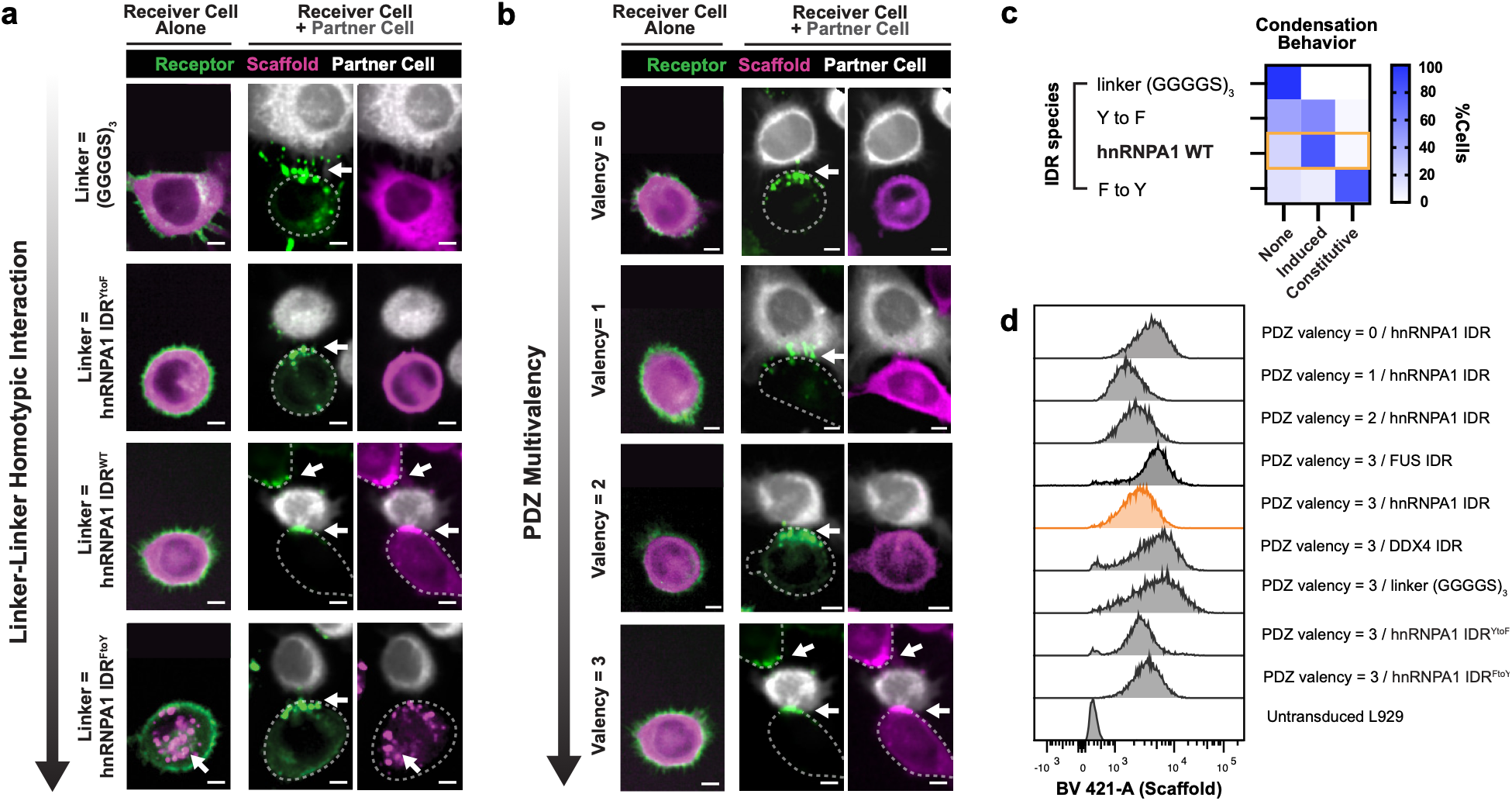
Characterization of PDZ scaffold variants and their condensation behaviors (linked to Main Figure 2b, c). Confocal microscopy images of receiver cells expressing PDZ scaffolds. Scale bar, 5 um. t = 2 h after co-culture with partner cells. GFP receptors are shown in green, PDZ scaffolds in magenta, and aGFP partner cells in white (labeled with CellTrace Far Red). a, Representative images of receiver cells expressing PDZ scaffolds bearing GS-rich linkers and mutants of hnRNPA1 IDR. b, Representative images of receiver cells expressing PDZ scaffolds with varied PDZ valency. c, Combinatorial matrix depicting scaffold condensation behaviors (%cells showing none, induced, or constitutive condensation) as a function of IDR/linkers. n = 60, 60, 60, and 51 for linker (GGGGS)_3_, hnRNPA1 IDR (Y to F mutant), hnRNPA1 IDR (WT), and hnRNPA1 IDR (F to Y mutant). d, Flow cytometry analysis of the PDZ scaffold variant expression levels in L929 fibroblast cells following cell sorting.

**Extended Data Figure 4.**
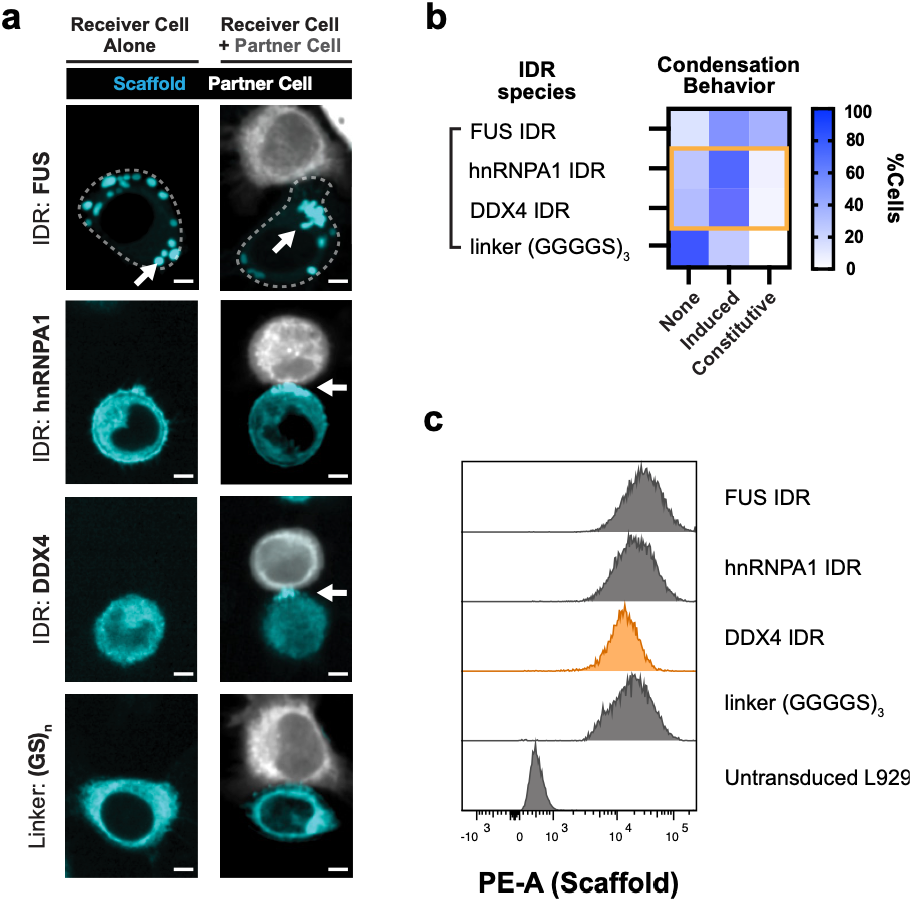
Distinct IDRs change condensation behaviors of receivers engineered with SYNZIP scaffolds (linked to Main Figure 2d, e). a, Representative confocal microscopy images of receiver cells engineered with SYNZIP scaffolds bearing different IDRs. Scaffold is shown in cyan, and partner cells in white (labelled with CellTrace Far Red). Scale bars, 5 µm. t = 2 h. b, Heatmap matrix depicting scaffold condensation state (%cells showing none, induced, or constitutive condensation) as a function of IDRs in receivers engineered with SYNZIP scaffolds. n = 69, 50, 67, and 49 for FUS IDR, hnRNPA1 IDR, DDX4 IDR, and linker (GGGGS)_3_, respectively. c, Flow cytometry analysis of SynZIP scaffolds showing matched expression following cell sorting.

**Extended Data Figure 5.**
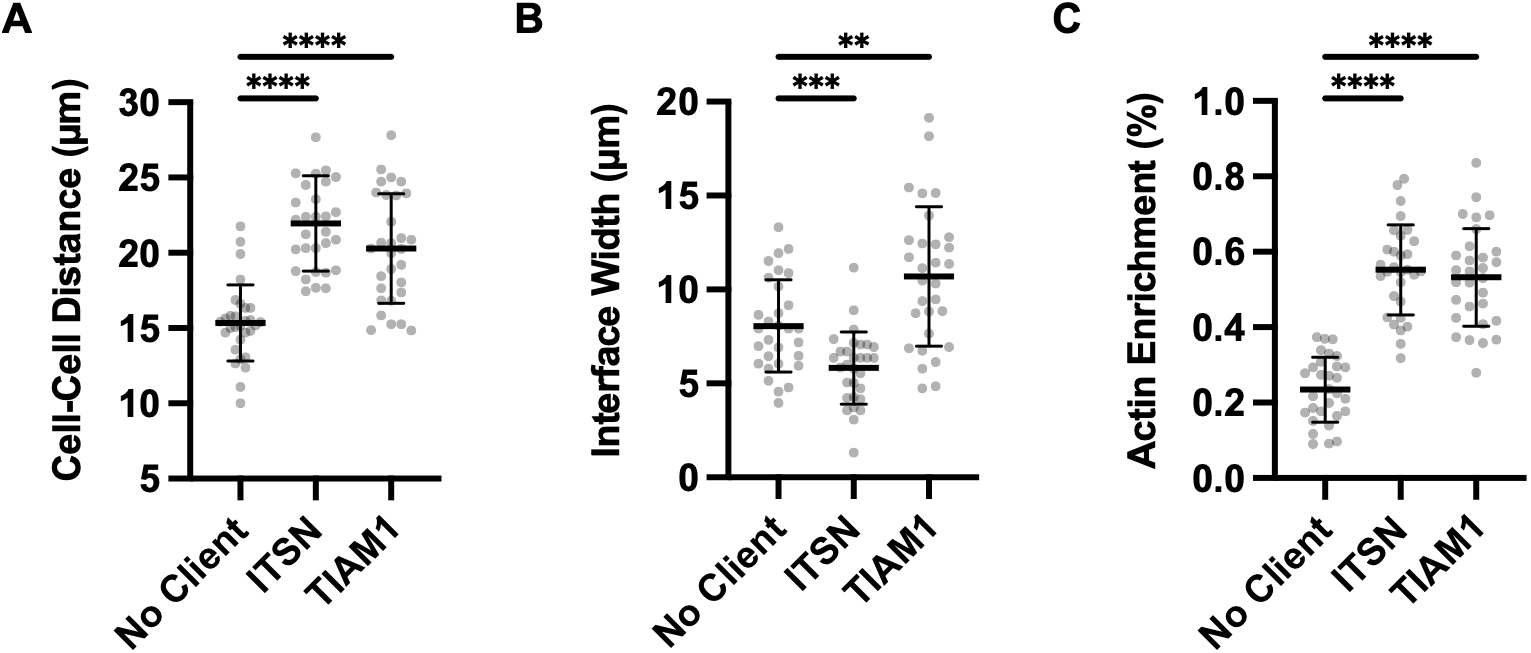
Quantification of actin-enriched interfaces induced by recruitment of Rho-family GTPase GEFs. L929 mouse fibroblasts expressing the GFP receptor and client-fused PDZ scaffold were fixed and analyzed 2 hours after contact with the cognate partner cell that expresses αGFP receptor and client-fused PDZ scaffold. Fixed cells were stained and quantified for a, cell–cell distance, b, interface width and c, relative actin enrichment at the cell–cell interface. Data are mean ± s.d.; n = 30 cell pairs per condition. Statistical significance was assessed using Brown–Forsythe and Welch ANOVA with the indicated post hoc comparisons. a, No client versus ITSN, P < 0.0001; no client versus TIAM1, P < 0.0001. b, No client versus ITSN, P = 0.0005; ITSN versus TIAM1, P = 0.0043. c, No client versus ITSN and no client versus TIAM1, P < 0.0001.

